# Presence of distinct operant phenotypes and transient withdrawal-induced escalation of operant ethanol intake in female rats

**DOI:** 10.1101/2024.09.05.611477

**Authors:** Joseph R Pitock, Shannon R Wheeler, Arleen Perez Ayala, Shikun Hou, Nathaly Arce Soto, Elizabeth J Glover

**Affiliations:** Center for Alcohol Research in Epigenetics, Department of Psychiatry, University of Illinois at Chicago, Chicago, IL, USA

**Keywords:** drinking, alcohol use disorder, sex differences, lickometer

## Abstract

Operant self-administration is frequently used to investigate the neurobiological mechanisms underlying alcohol seeking and drinking and to test the efficacy of drugs under development for the treatment of alcohol use disorder (AUD). Although widely used by the research community, there is a paucity of operant ethanol self-administration studies that include female subjects. The current study characterizes home cage drinking and operant ethanol self-administration in female Sprague Dawley, Long Evans, and Wistar rats. Rats underwent three weeks of intermittent-access two-bottle choice home cage drinking before being trained to lever press for ethanol in standard operant chambers equipped with contact lickometers. After capturing baseline operant performance, rats were chronically exposed to control or ethanol liquid diet using the Lieber-DeCarli method. Operant ethanol self-administration was re-evaluated after chronic liquid diet exposure to determine whether female rats exhibit similar withdrawal-induced escalation of ethanol intake as is regularly observed in male rats. Our findings reveal the presence of three distinct operant phenotypes (Drinker, Responder, Nonresponder), the prevalence of which within each strain is strikingly similar to our previous observations in males. Within a given phenotype, rats of each strain performed similarly during operant testing. Ethanol intake during home cage drinking was unable to predict future operant phenotype. Relative to controls, Drinkers chronically exposed to ethanol liquid diet exhibited a significant, but transient, escalation in consummatory, but not appetitive, responding during acute withdrawal. Collectively, these data closely parallel many of our previous observations in males while also highlighting potential sex differences in drinking strategies following dependence. The presence of the Responder phenotype reinforces the importance of using direct measures of ethanol consumption. Our findings provide new insight into similarities and differences in operant ethanol self-administration between males and females and emphasize the importance of including females in future studies of ethanol drinking and dependence.

## 1. Introduction

Alcohol use disorder (AUD) is a notable and prevalent health concern that has seen a rise in diagnoses in the US [1]. AUD severity is directly linked to morbidity and mortality with greater severity associated with increased risk for hospitalization [2] and serious negative health outcomes including learning and memory problems, weakened immune system, liver disease, heart disease, and stroke [3]. Taken together, this work highlights the public health crisis posed by alcohol misuse.

Operant ethanol self-administration is a well-known paradigm that is used preclinically to interrogate the neurobiological mechanisms underlying drinking behavior and to test potential interventions for AUD. For example, this paradigm is routinely used to investigate the mechanisms responsible for escalation of ethanol intake – a model of uncontrolled drinking. Using this approach, previous work has shown that ethanol dependent rats exhibit increased lever pressing for ethanol relative to pre-dependence levels [4–6]. The success of this model is further demonstrated in the pivotal role that operant ethanol self-administration played in the development of naltrexone for the treatment of AUD [7–9] and its continued use to test newly developed drugs with therapeutic potential [10–13].

Operant paradigms offer greater face validity over home cage drinking methods by requiring that individuals work to acquire alcohol similar to real-world circumstances (e.g., going to the bar/store to purchase liquor). Perhaps as a result, the majority of studies using this approach tend to report ethanol intake in terms of the work required (i.e., active lever presses, number of reinforcers earned) with consumption inferred based on appetitive responses or reinforcer delivery. Although common, this approach has increasingly been called into question as more sophisticated methods that use direct measurements of consumption show that reinforcer delivery is not always directly linked with consumption [14–16]. This includes our own work [17], which measured operant performance in three rat strains commonly used in alcohol research. Using a contact lickometer-equipped system, our work uncovered three distinct “operant phenotypes” (Drinker, Responder, Nonresponder) with differing prevalence across strains. These phenotypes provide insight into the potential influence of strain on the successful acquisition of operant self-administration. They further reveal that a significant number of rats refrain from drinking despite regularly completing the necessary operant response requirement to gain access to ethanol – behavior that would go undetected in the absence of direct measures of consumption.

Historically, the vast majority of operant ethanol self-administration studies, including our own, have been restricted to male subjects [17, 18]. This sentiment is supported by a survey of journal articles published in 2009, which found that bias toward male subjects was most prominent in the field of neuroscience than any other biological discipline [19]. A similar, albeit informal, survey of Web of Science found that only 7% of articles returned with the key terms “alcohol dependence” or “ethanol dependence” and “rat” also included the term “female” [20]. This lack of female inclusion persists despite well-documented sex and gender differences in alcohol consumption as well as AUD prevalence and prognosis. Thus, while men consume more alcohol on average than women [21] and are more likely to be diagnosed with AUD [1], this gender gap has narrowed in recent years [22]. Moreover, data suggests that women may progress from casual drinking to AUD diagnosis more rapidly [23] and experience higher rates of morbidity [24–26] than men. Therefore, inclusion of female, as well as male, subjects in drinking studies is crucial in order to identify similarities and differences in the neurobiological mechanisms underlying drinking behavior between sexes.

The present study builds upon our prior work in males to explore strain differences in home cage and operant ethanol self-administration, as well as the potential for withdrawal-induced escalation of intake in female rats. Our findings uncover the presence of the same three operant phenotypes previously characterized in males [17] in females and replicate the same difference in prevalence of each phenotype across the three strains most frequently used in alcohol research. This includes a subset of female rats that do not consume ethanol upon completion of the operant response requirement. We further show that withdrawal-induced escalation of operant ethanol self-administration is transient in females, unlike the persistent escalation previously observed in males.

## 2. Materials and Methods

### 2.1 Animals

Adult female Long-Evans (n = 16; Envigo, Indianapolis, IN), Sprague Dawley (n = 16; Envigo), and Wistar (n = 18; Charles River, Wilmington, MA) rats were P57-60 upon arrival. Rats were single-housed in standard polycarbonate cages in a room operating on a reverse 12:12 light-dark cycle (lights on at 22:00) and allowed to acclimate to the vivarium for at least one week prior to beginning the experiment. Rats had *ad libitum* access to standard chow (Teklad 7912, Envigo) and water unless otherwise stated. All experimental procedures adhered to the University of Illinois at Chicago Institutional Animal Care and Use Committee and followed the NIH Guidelines for the Care and Use of Laboratory Animals.

### 2.2 Intermittent-access two-bottle choice home cage drinking

Intermittent-access two-bottle choice home cage drinking was performed according to previously published procedures [17, 18]. Briefly, over the course of three weeks, animals were provided with 24-hour access to two bottles. On Monday, Wednesday, and Friday, 30 minutes into their dark cycle, rats were provided access to one bottle containing water and another containing ethanol (20% v/v). The ethanol bottle was placed on alternating sides on each day to avoid side preference. On all other days, animals were provided with two bottles containing water. Consumption was measured by calculating change in bottle weight at 30 minutes and 24 hours after the onset of the drinking session. Weight loss from water and ethanol bottles placed on an empty cage was subtracted from bottle weight change calculations made for each rat on each day to correct for fluid loss due to spillage.

### 2.3 Operant self-administration

Following three weeks of home cage drinking, rats were trained to operantly self-administer ethanol according to previously published procedures [17] using standard operant boxes housed in sound attenuating chambers (Med Associates, St Albans, VT). Each box was affixed with a house light, two retractable levers, a cue light above each lever, and an opening that allowed access to a retractable lickometer-equipped sipper tube containing ethanol (20% v/v). This allowed for direct measurement of appetitive (active lever presses and reinforcers delivered) and consummatory (licks and ethanol intake in g/kg) measures during each session. Rats were trained to lever press for access to ethanol during sessions that progressively decreased in duration in parallel with an increase in the response requirement. Rats transitioned to each subsequent training phase after stable responding had been maintained (∼10-15 sessions). During the first phase, rats were trained to lever press in 60 min sessions on an FR1 schedule of reinforcement, during which one active lever press resulted in retraction of the active and inactive levers, illumination of the cue light above the active lever, and 15 seconds of access to the ethanol-containing sipper tube. Inactive lever presses were recorded but had no programmed response. Next, session duration was decreased to 30 min. After stable responding had been achieved, the response requirement increased to FR3. Then, session duration was further shortened to 15 minutes. Ethanol intake during each session was measured by calculating change in bottle weight measured before and after each session normalized for average spillage calculated based on bottle weight change from sessions with zero licks (0.05 g). Intake was calculated by body weight taken daily after rats completed each session.

### 2.4 Lieber DeCarli liquid diet

After acquiring baseline levels of operant ethanol self-administration, rats were either rendered dependent or maintained on control diet using the Lieber DeCarli liquid diet procedure following previously published procedures [17, 27]. During this time, liquid diet was the sole source of fluid and nutrition for all rats. For the first three days of the procedure, all rats, regardless of group assignment, received control liquid diet after which rats assigned to the ethanol group received diet with increasing concentrations of ethanol over the course of seven days (from 1.8% to 9%).

Rats in the ethanol group were then maintained on 9% ethanol for 15 days. Rats assigned to the control group received control liquid diet for the duration of the experiment in a manner that was yoked to consumption exhibited by rats in the ethanol group.

### 2.5 Withdrawal-induced escalation

Rats resumed operant testing on an FR3 schedule of reinforcement during 15 min sessions upon completion of the liquid diet procedure. Testing was performed during acute withdrawal according to previously published work [17]. For three weeks, on Monday and Thursday, rats in the ethanol group received control diet instead of their standard ethanol diet and operant performance was tested the following day approximately 24 hours later for a total of six acute withdrawal operant test sessions. Rats in the ethanol group received ethanol liquid diet upon return to the home cage after completion of each operant session. Control rats continued to receive control liquid diet yoked to the ethanol group during this time.

### 2.6 Statistical analysis

One-way, two-way, and multifactorial ANOVAs were used to analyze strain and phenotype differences in home cage and operant performance as specified below. Mixed-effects ANOVAs were substituted for the aforementioned analyses when missing values were present in the data set. When included as a factor, session was defined as a repeated measure (RM). Baseline operant performance was calculated from the average of the last five operant sessions before undergoing the liquid diet procedure. Spearman’s correlations were used to determine the relationship between appetitive and consummatory responding at baseline as well as the relationship between home cage and operant drinking. All analyses were performed using Prism 10 (GraphPad, Boston, MA). Significance was set at p≤0.05 and all data are presented as mean ± SEM.

## 3. Results

### 3.1 Ethanol intake is similar in female rats of all three strains during home cage drinking

Intermittent-access two-bottle choice consistently produces ethanol intake in males that plateaus after a short escalation phase at the onset of the drinking paradigm [17, 28]. In contrast to this, no clear pattern of escalation was observed in females of all three strains in the present study. Instead, female rats drank relatively consistent levels of ethanol throughout the home cage drinking period. In addition, intake was similar across all three strains. This was confirmed using a mixed-effects ANOVA to examine 24 hr intake, which found no significant main effect of either drinking day [F (4.591, 215.2) = 2.097; p=0.0728] or strain [F (2, 47) = 1.497; p=0.2342] or a significant interaction between factors [F (16, 375) = 1.267; p=0.2154] (**Figure 1A**). Although a similar analysis of ethanol preference during 24 hr intake revealed a significant interaction between drinking day and strain [F (16, 376) = 1.9; p=0.0164], posthoc comparisons failed to uncover significant strain differences on any given drinking day (**Figure 1C**). In the absence of a significant main effect of strain [F (2, 47) = 0.68; p=0.5121], these data further suggest that average daily intake is similar across strains of female rats.

**Figure 1.**
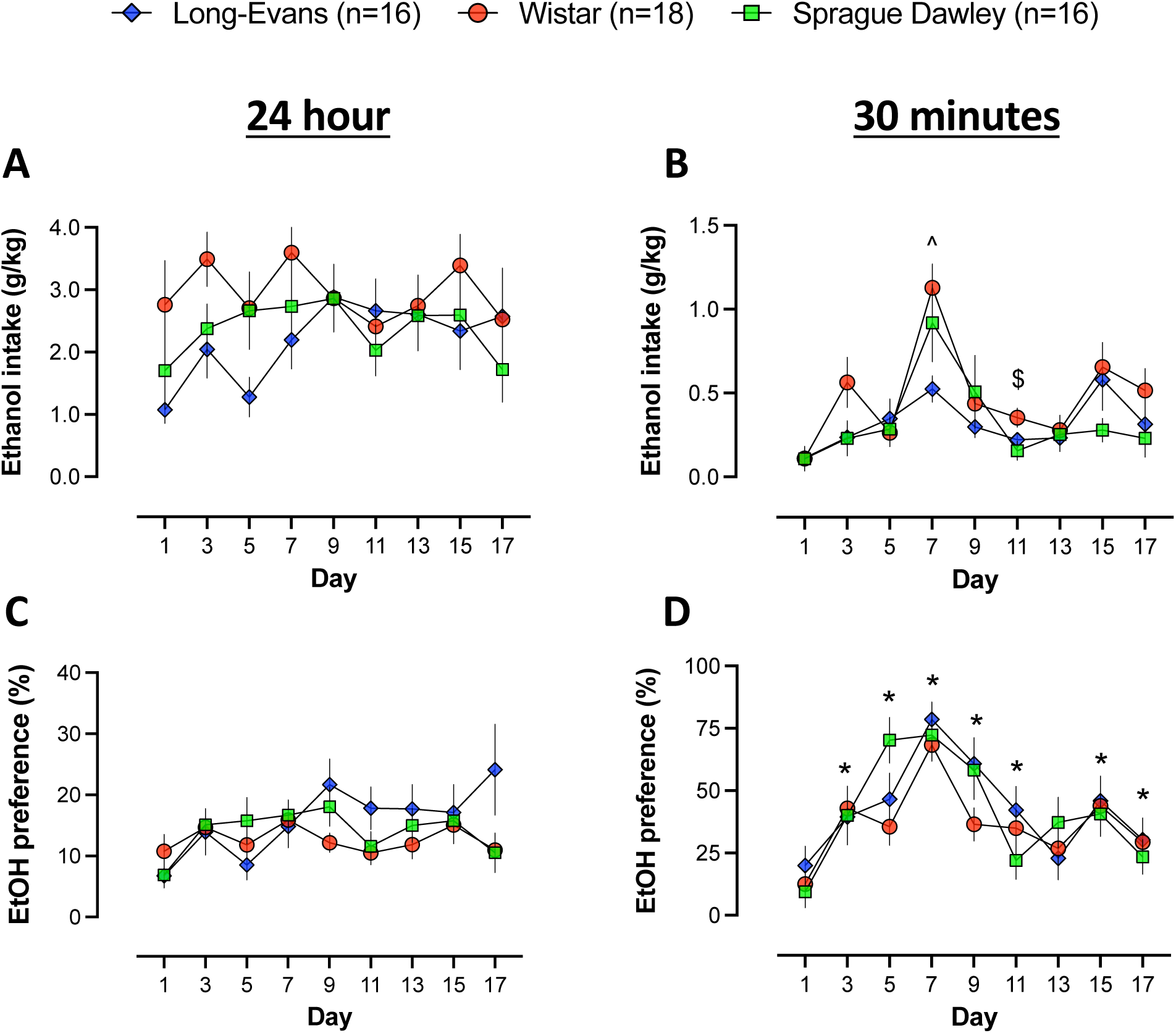
Few strain differences in intermittent-access two-bottle choice ethanol intake or preference in female rats. **(A)** Average daily ethanol intake does not differ across drinking days or between strains. **(B)** Average 30 min ethanol intake was similar across strains on the majority of drinking days with Long-Evans and Sprague Dawley rats drinking significantly less ethanol than Wistars on days 7 and 11, respectively. All three strains exhibited a similar degree of ethanol preference at both 24 hr **(C)** and 30 min **(D)** although preference at 30 min varied significantly from day 1 on subsequent drinking days. ^p<0.05 Long-Evans compared to Wistar; ^$^p<0.05 Sprague Dawley compared to Wistar; *p<0.05 compared to day 1.

Somewhat greater variability was observed between strains when examining ethanol intake during the first 30 min of bottle access. This was confirmed using a two-way RM ANOVA, which uncovered a significant main effect of drinking day [F (5.347, 251.3) = 13.12; p<0.0001] and an interaction between drinking day and strain [F (16, 376) = 1.746; p=0.0369] in the absence of a main effect of strain [F (2, 47) = 2.154; p=0.1273]. Tukey corrected post-hoc comparisons found that Long-Evans and Sprague Dawley rats drank significantly less than Wistars on days 7 (p=0.0027) and 11 (p=0.0474), respectively (**Figure 1B**). Nevertheless, no strain differences were observed when examining ethanol preference at the 30-min time point. This was evident using a mixed-effects ANOVA, which did not identify significant effects of strain [F (2, 47) = 0.57; p=0.5719] or an interaction between strain and drinking day [F (16, 375) = 1.3; p=0.1876]. This analysis did, however, uncover a significant main effect of drinking day [F (5.4, 255) = 15; p<0.0001] with Sidak’s multiple comparisons finding that preference for ethanol was significantly higher on all subsequent drinking days relative to the first day except for day 13 (all p values < 0.05; **Figure 1D**).

### 3.2 Strain differences in operant phenotypes in females parallel previous observations in males

Our previous work identified the presence of three distinct operant phenotypes in male rats with representation of each phenotype varying across rat strains [17]. To explore whether the same phenotypes were also present in females we categorized rats based on our previously published criteria: 1) Drinkers are rats that, on average, gain access to >2 reinforcers and consume >0.1 g/kg/session, during baseline operant sessions; 2) Responders are rats that, on average, gain access to >2 reinforcers but consume <0.1 g/kg/session, during baseline operant sessions; 3) Nonresponders are rats whose responding is associated with <2 reinforcers and <0.1 g/kg intake per session on average during baseline operant sessions [17] (**Figure 2A-D**). This assessment confirmed that, similar to our previously published work in males, all three operant phenotypes are present in female rats of all three strains. Of note, Long-Evans rats exhibited the highest proportion of Drinkers (44%) and lowest proportion of Responders (19%) (**Figure 2E**). In contrast, Responders constituted the majority phenotype in both Wistar (**Figure 2F**) and Sprague Dawley (**Figure 2G**) strains. Altogether, these data closely parallel our observations in male rats.

**Figure 2.**
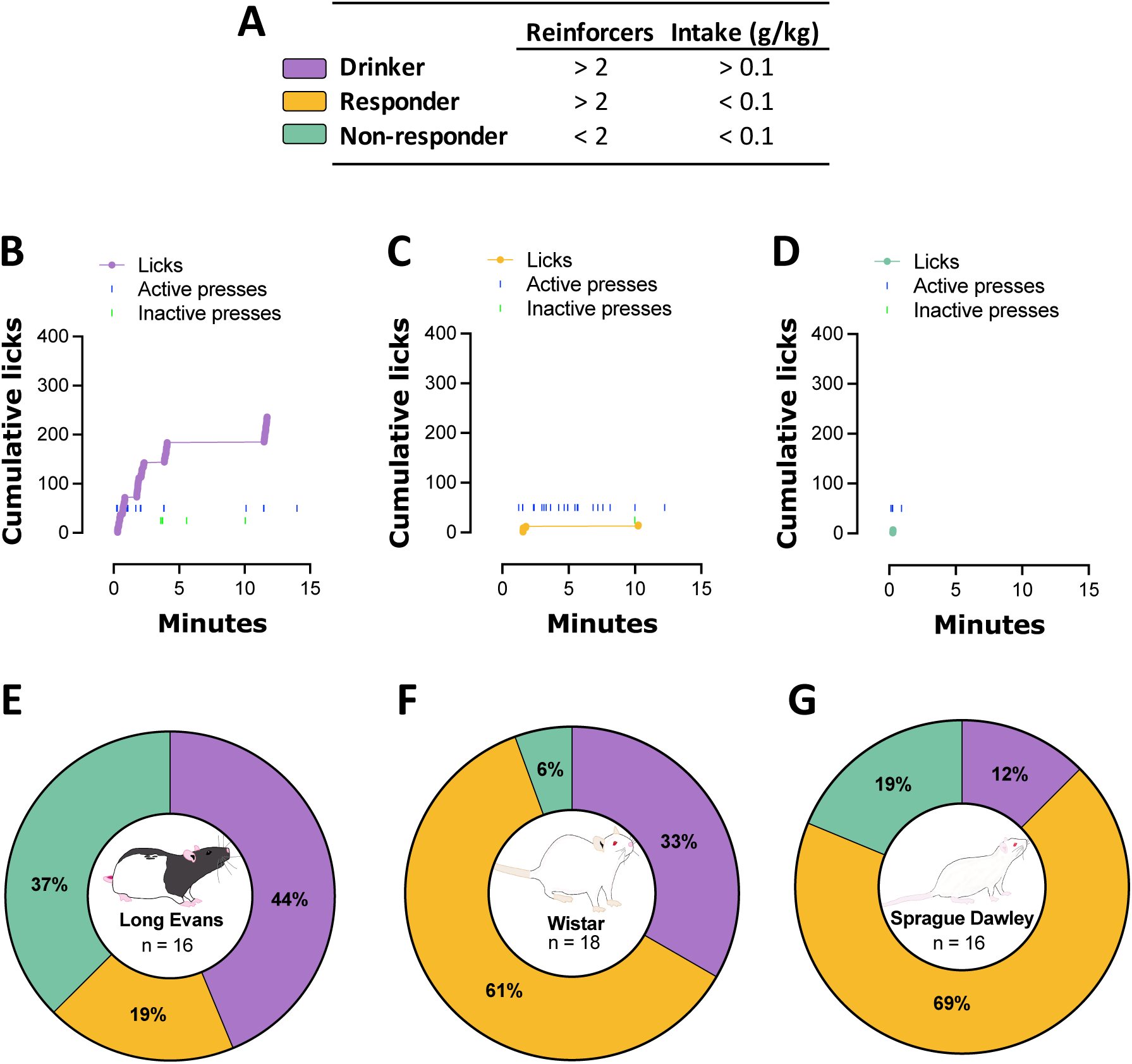
Operant phenotypes are differentially represented across three strains in female rats. **(A)** Criteria used for phenotypic classification (based on Patwell et al., 2021). Representative cumulative records of a Drinker **(B)**, Responder **(C)**, and Non-responder **(D)** reveal distinct appetitive and consummatory patterns of behavior within each phenotype. While all three phenotypes were present in all three rat strains, the greatest proportion of Drinkers was present in Long-Evans rats **(E)** with the majority of the remainder of this strain falling into the Non-responder category and relatively few rats classified as Responders. A smaller proportion of Wistar rats **(F)** met the criteria for Drinkers. Instead, the majority in this strain were classified as Responders. A similarly high degree of Responders was present in the Sprague Dawley strain **(G)** with relatively fewer rats meeting Drinker criteria.

### 3.3 Baseline operant performance is similar across strains within each phenotype

To evaluate potential strain differences in operant performance, we next analyzed appetitive and consummatory behaviors at baseline within Drinker and Responder phenotypes. Of note, Sprague Dawley rats (n=2) and Long-Evans rats (n=3) were omitted from the analysis of Drinkers and Responders, respectively, because their low representation resulted in insufficient power for statistical analysis of these strains within these phenotypes. A multifactorial ANOVA comparing lever pressing across strains and sessions in Drinkers confirmed that both Wistar and Long-Evans rats successfully discriminated between the active and inactive levers as evidenced by a significant main effect of lever [F (1.000, 11.00) = 129.1; p<0.0001]. Therefore, subsequent analysis was limited to active lever pressing. A two-way RM ANOVA of active presses found no significant effect of strain [F (1, 11) = 0.09817; p=0.7599] or session [F (2.165, 23.82) = 0.1274; p=0.8952] but did uncover a significant interaction between session and strain [F (4, 44) = 2.703; p=0.0425]. However, post-hoc comparisons using a Sidak correction failed to uncover significant between-strain differences in active lever presses during any given operant session (all p values 0.5; **Figure 3A**) suggesting that only marginal differences drove the observed significant interaction between factors. Similar results were observed when performing the same analysis of average number of reinforcers (i.e., sipper access). Here, a two-way RM ANOVA found a significant interaction between strain and operant session [F (4, 44) = 2.571; p=0.0509] in the absence of significant main effects of strain [F (1, 11) = 0.04141; p=0.8425] or session [F (2.205, 24.25) = 0.1592; p=0.8724]. Once again post-hoc comparisons failed to uncover significant differences in reinforcers earned between strains during any given operant session (all p values 0.5; **Figure 3C**). Consummatory behaviors largely paralleled appetitive responding in Drinkers. A two-way RM ANOVA of average licks revealed a significant strain by session interaction [F (4, 44) = 2.764; p=0.0391] in the absence of main effects [strain: F (1, 11) = 0.2249; p=0.6446; session: F (1.781, 19.59) = 0.5479; p=0.5669]. Once again, post-hoc comparisons did not uncover significant between-strain differences in average number of licks during any given operant session (all p values > 0.1; **Figure 3E**). Ethanol intake was also similar between strains with a two-way RM ANOVA finding no significant main effects [strain: F (1, 11) = 1.685; p=0.2209; session: F (2.385, 25.64) = 0.1095; p=0.9249] or strain by session interaction [F (4, 43) = 2.510; p=0.0557] (**Figure 3G**). Altogether, these data reveal an absence of strain differences in either appetitive or consummatory behaviors in Drinkers trained to operantly self-administer ethanol.

**Figure 3.**
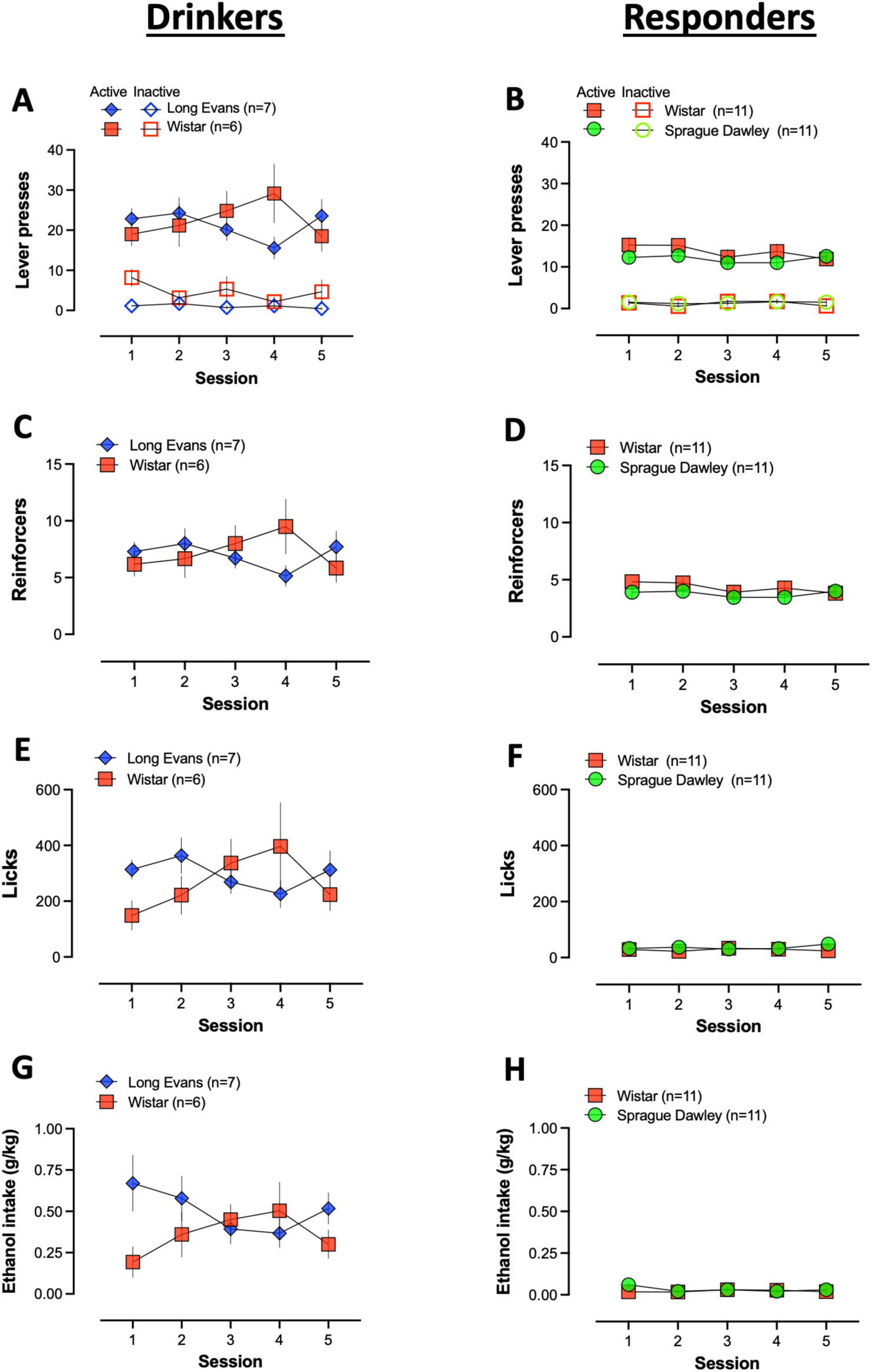
Similar behavioral performance between strains within phenotypes. Rats of different strains perform similarly during operant sessions within Drinker **(A,C,E,G)** and Responder **(B,D,F,H)** phenotypes. Both Drinkers **(A)** and Responders **(B)** exhibit robust active lever pressing during sessions and display strong discrimination between the active and inactive levers. Despite similar levels of access to the reinforcer **(C, D)**, Drinkers exhibit far greater number of licks **(E)** and intake **(G)** than Responders who perform few licks **(F)** and have intake **(H)** that does not exceed 0.1 g/kg.

Similar to Drinkers, rats in the Responder phenotype successfully discriminated between active and inactive levers. This was evident using a multifactorial ANOVA, which found a significant main effect of lever [F (0.6643, 13.29) = 137.8; p<0.0001]. Two-way RM ANOVAs were subsequently used to assess strain differences in appetitive responding within Responder rats. These analyses found no significant effects of strain [lever press: F (1, 20) = 0.7768; p=0.3886; reinforcer: F (1, 20) = 0.6178; p=0.4411], session [lever press: F (2.874, 57.48) = 2.624; p=0.0614; reinforcer: F (3.035, 60.70) = 2.294; p=0.0862], or strain by session interaction [lever press: F (4, 80) = 1.435; p=0.2301; reinforcer: F (4, 80) = 1.154; p=0.3374] for either average active lever presses (**Figure 3B**) or reinforcers earned (**Figure 3D**). Although consumption was, by definition, extremely low in Responders, we applied the same analysis as was used in Drinkers to explore the possibility that different strains of Responders exhibited different degrees of consumption during operant sessions. A two-way RM ANOVA of average licks across sessions found no significant main effects of strain [F (1, 20) = 1.140; p=0.2984] or session [F (2.233, 44.66) = 0.6288; p=0.5548]. This analysis did, however, uncover a significant strain by session interaction [F (4, 80) = 2.965; p=0.0244], although post-hoc comparisons failed to identify significant between-strain differences in average licks performed on any given session (all p values > 0.3; **Figure 3F**). No significant main effects [strain: F (1, 20) = 1.960; p=0.1769; session: F (2.593, 51.86) = 1.420; p=0.2496] or interaction between factors [F (4, 80) = 2.156; p=0.0815] were observed with regard to ethanol intake (**Figure 3H**). These data show that while Wistar and Sprague Dawley Responders alike are able to successfully gain reinforcer access, they decline to consume substantive quantities of ethanol.

Spearman’s correlations were used to more directly interrogate the relationship between appetitive and consummatory behaviors within each phenotype. As expected, this analysis revealed as significant positive relationship between active pressing and reinforcers earned as well as licks and intake in both Drinkers (appetitive: r=0.996, p<0.001; consummatory: r=0.740, p<0.01) and Responders (appetitive: r=0.995, p<0.001; consummatory: r=0.723, p<0.001). In Drinkers, a significant positive correlation was also present between the number of active lever presses and licks (r=0.797, p<0.01) and intake (r=0.819, p<0.001). A similar significant positive relationship was observed between reinforcers earned and licks (r=0.814, p<0.001) and intake (r=0.812, p<0.001) in this phenotype (**Figure 4A,C**). In contrast, consummatory behaviors were not significantly correlated with appetitive responding in the Responder phenotype (active press x licks: r=0.085, p=0.685; active press x intake r=0.013, p=0.952; reinforcers x licks: r=0.059, p=0.778; reinforcers x intake: r=0.004, p=0.984) (**Figure 4B,D**). Taken together, these data show that appetitive measures are not predictive of consumption in all rats.

**Figure 4.**
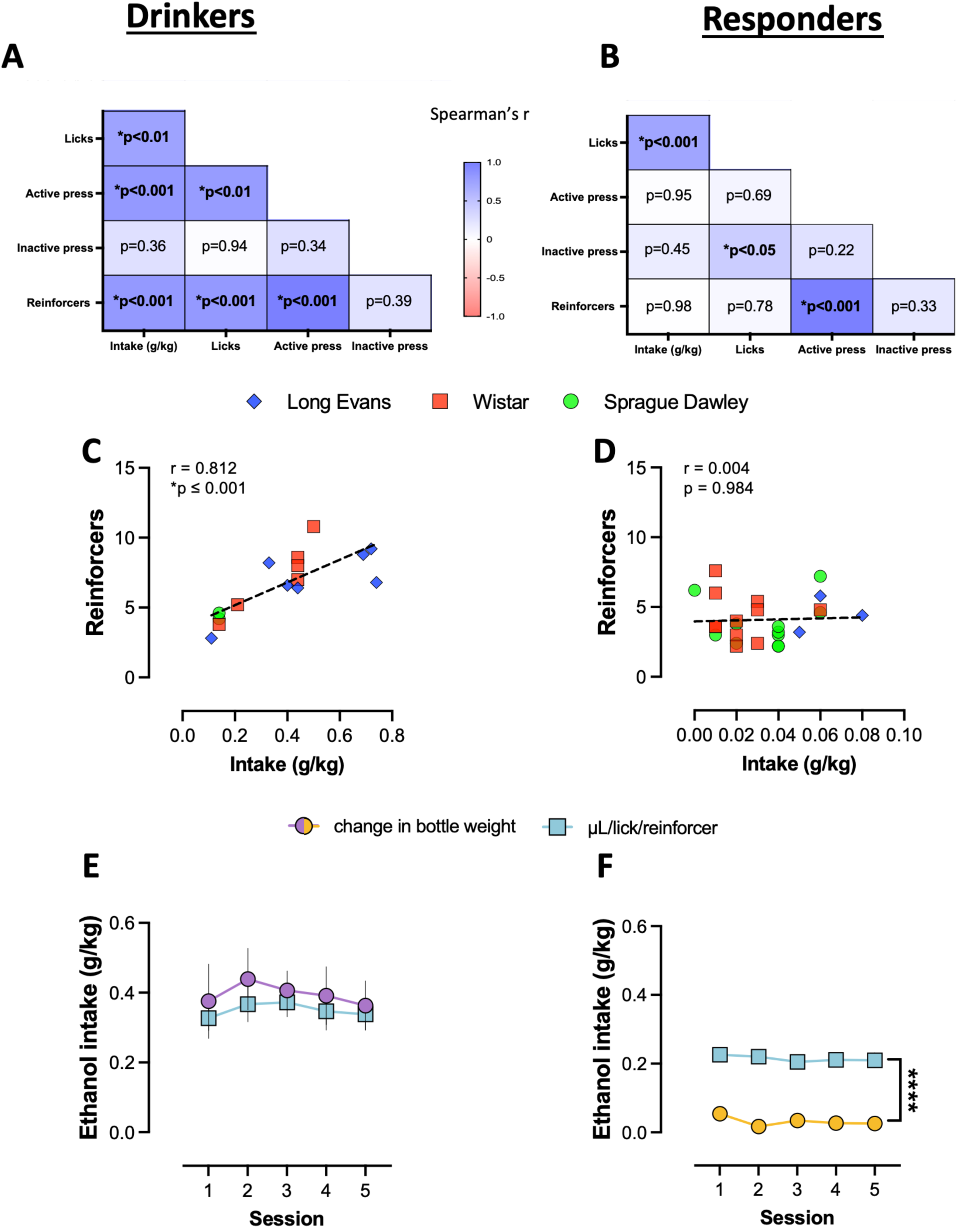
Appetitive responding correlates with consummatory behaviors only in the Drinker phenotype. **(A)** In Drinkers, measures of appetitive responding are significantly positively correlated with measures of consumption. **(B)** No significant relationship is observed between these same measures in the Responder phenotype. This is further highlighted when plotting the relationship between reinforcers earned and ethanol intake in Drinker **(C)** and Responder **(D)** phenotypes. Intake inferred from reinforcer delivery (μL/lick/reinforcer) is similar to intake calculated based on change in bottle weight in Drinkers **(E)** but not Responders **(F)**. ****p<0.0001.

To further emphasize the risk of generating misleading results when appetitive measures are used to infer consumption, we next compared intake calculated based on change in bottle weight with intake estimated from reinforcer delivery. Two constants were developed from data generated in Drinkers to calculate estimated intake: 1) The average number of licks performed per reinforcer access was calculated by dividing total licks by total reinforcers during each baseline session per Drinker (35.76 ± 1.71); 2) The average volume of ethanol consumed per lick (μL) was calculated by dividing the change in bottle volume (mL) by total licks during each baseline session (2.56 ± 0.15). Using these constants, we estimated that each reinforcer access period is associated with consumption of 91.54 μL ethanol. Interestingly, this value is remarkably close to the 100 μL volume of ethanol commonly delivered upon completion of the operant response requirement in studies using the liquid receptacle system (e.g., [4, 5, 15, 29, 30]). As shown in **Figure 4E**, a two-way RM ANOVA in Drinkers found no significant differences in intake calculated based on change in bottle weight versus that inferred by number of reinforcers [Intake measure: F (1, 28) = 0.4589; p=0.5037; Session: F (2.554, 70.86) = 0.4095; p=0.7148; Intake measure by Session: F (4, 111) = 0.05527; p=0.9942]. In contrast, the same analysis in Responders (**Figure 4F**) revealed a significant main effect of intake measure [F (1, 48) = 112.9; p<0.0001] with estimates of over 0.20 g/kg for these rats when intake was inferred from reinforcer delivery. No significant main effect of session [F (3.342, 160.4) = 1.853, p=0.1331] or intake measure by session interaction [F (4, 192) = 0.8888, p=0.4717] was observed. These data further emphasize the importance of using direct measures to accurately determine ethanol consumption during operant self-administration sessions.

### 3.4 Operant phenotypes are indistinguishable during home cage drinking

Previous work has suggested that high levels of ethanol intake in the home cage may be indicative of high levels of intake during operant self-administration [31]. To explore this, we next examined whether indices of consumption during intermittent-access two-bottle choice home cage drinking differed across operant phenotypes. A one-way ANOVA showed that average intake during the first 30 min of the drinking session was similar across all three phenotypes [F (2, 47) = 0.05567; p=0.9459] (**Figure 5A**). Average 24 hr ethanol intake was also similar between future Drinkers, Responders, and Nonresponders [F (2, 47) = 0.007024; p=0.9930] (**Figure 5B**). Preference for ethanol was also not significantly different between phenotypes at either the 30-min [F (2, 47) = 0.4611; p=0.6334] or 24-hr [F (2, 47) = 0.2226; p=0.8012] time points (**Figure 5C-D**). In addition, Spearman’s correlations found no significant relationship between home cage ethanol intake at 30 min or 24 hr and baseline operant intake (**Figure 5E-F**). This was true for both Drinkers (30 min: r=0.2297, p=0.4073; 24 hr: r=-0.2351, p=0.3961) and Responders (30 min: r=0.09059, p=0.6962; 24 hr: r=0.09701, p=0.6757). These data show that the magnitude of ethanol intake during home cage drinking was not predictive of the magnitude of intake during operant ethanol self-administration.

**Figure 5.**
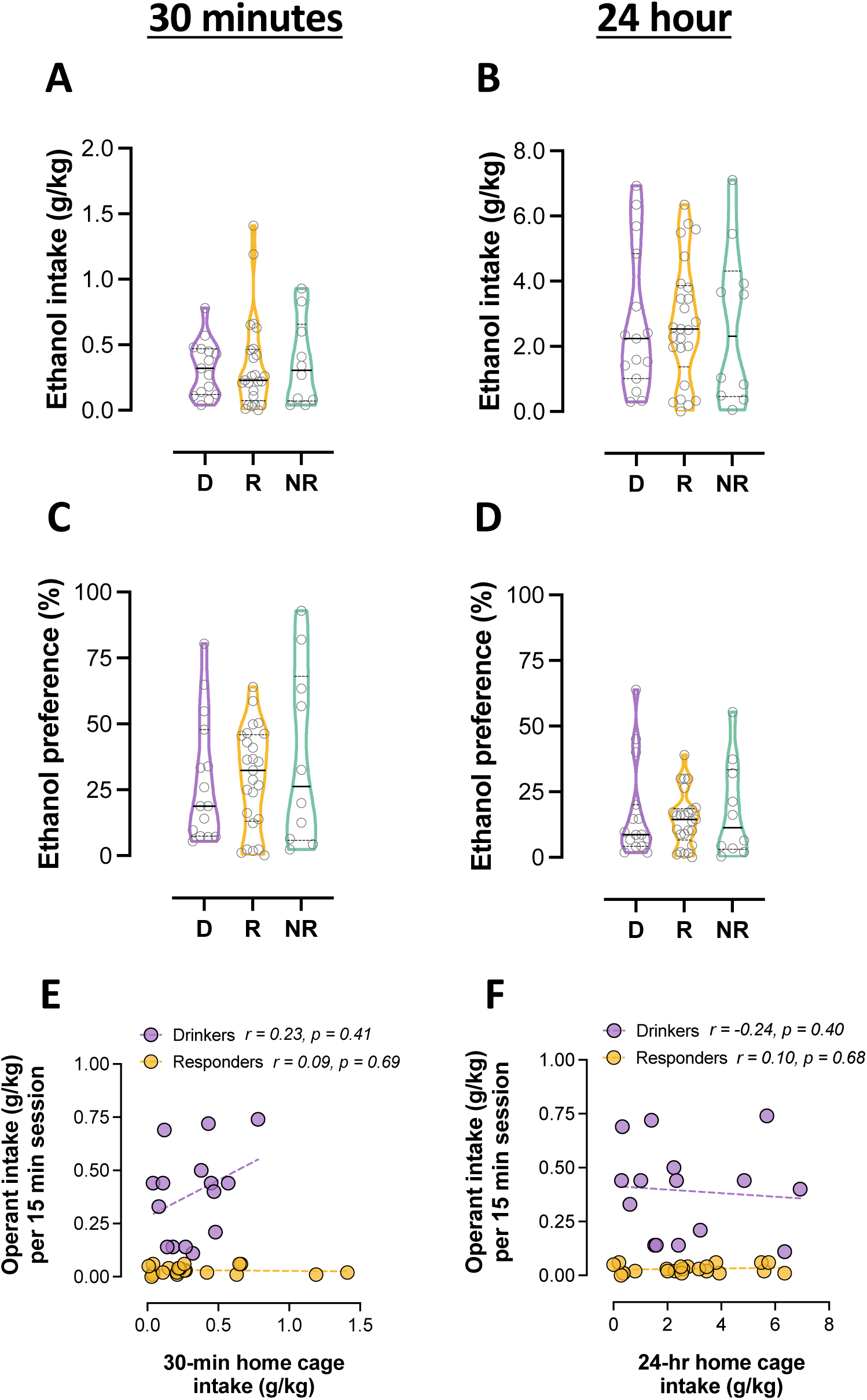
No relationship between home cage and operant drinking. Rats of all three phenotypes exhibit similar levels of home cage ethanol intake at 30 min **(A)** and 24 hr **(B)**. They also exhibit similar degrees of ethanol preference during home cage drinking at both 30 min **(C)** and 24 hr **(D)**. No significant correlations are observed when examining the relationship between ethanol intake during operant sessions and intake at either 30 min **(E)** or 24 hr **(F)** during home cage drinking.

### 3.5 Withdrawal-induced escalation of operant ethanol intake is transient in female Drinkers

In our prior work, we showed that male Drinkers exhibited a significant escalation in both appetitive and consummatory behaviors during acute withdrawal following chronic ethanol liquid diet exposure [17]. To explore whether withdrawal produces a similar effect in females, we compared operant performance at baseline to performance during repeated episodes of acute withdrawal. Based on the findings above showing no strain differences in operant performance at baseline in combination with reduced representation of each strain as a result of splitting into control and ethanol liquid diet groups, data were collapsed by strain for these analyses. Two-way RM ANOVAs were first used to measure the effect of withdrawal on measures of appetitive responding in Drinkers. No significant main effects of liquid diet exposure [F (1, 13) = 0.07661; p=0.7863] or time [F (3.846, 50.00) = 1.636; p=0.1818] or interaction between the two factors [F (6, 78) = 1.459; p=0.2032] was found for average number of active lever presses per session (**Figure 6A**). The same analysis found no significant main effects [diet: F (1, 13) = 0.02348; p=0.8806; time: F (3.889, 50.56) = 1.659; p=0.1755] or interaction [F (6, 78) = 1.451; p=0.2064] with respect to reinforcer access (**Figure 6C**). In contrast to appetitive behavior, the number of licks per session increased during withdrawal relative to baseline in female rats exposed to ethanol but not control liquid diet (**Figure 6E**). However, the interaction between time and liquid diet approached, but did not reach, statistical significance [F (6, 78) = 1.971; p=0.0799] using a two-way RM ANOVA. No significant main effects of either liquid diet [F (1, 13) = 1.819; p=0.2004] or time [F (3.746, 48.70) = 2.078; p=0.1021] were observed in this analysis. A two-way RM ANOVA of ethanol intake found no main effect of diet [F (1, 13) = 1.748; p=0.2089] or a significant interaction between diet and time [F (6, 78) = 1.729; p=0.1253]. This analysis did uncover a significant main effect of time [F (3.421, 44.47) = 2.820; p=0.0432]. However, posthoc comparisons using a Dunnett correction failed to uncover any sessions during acute withdrawal for which ethanol intake was significantly different than intake during baseline (all p values > 0.10; **Figure 6G**).

**Figure 6.**
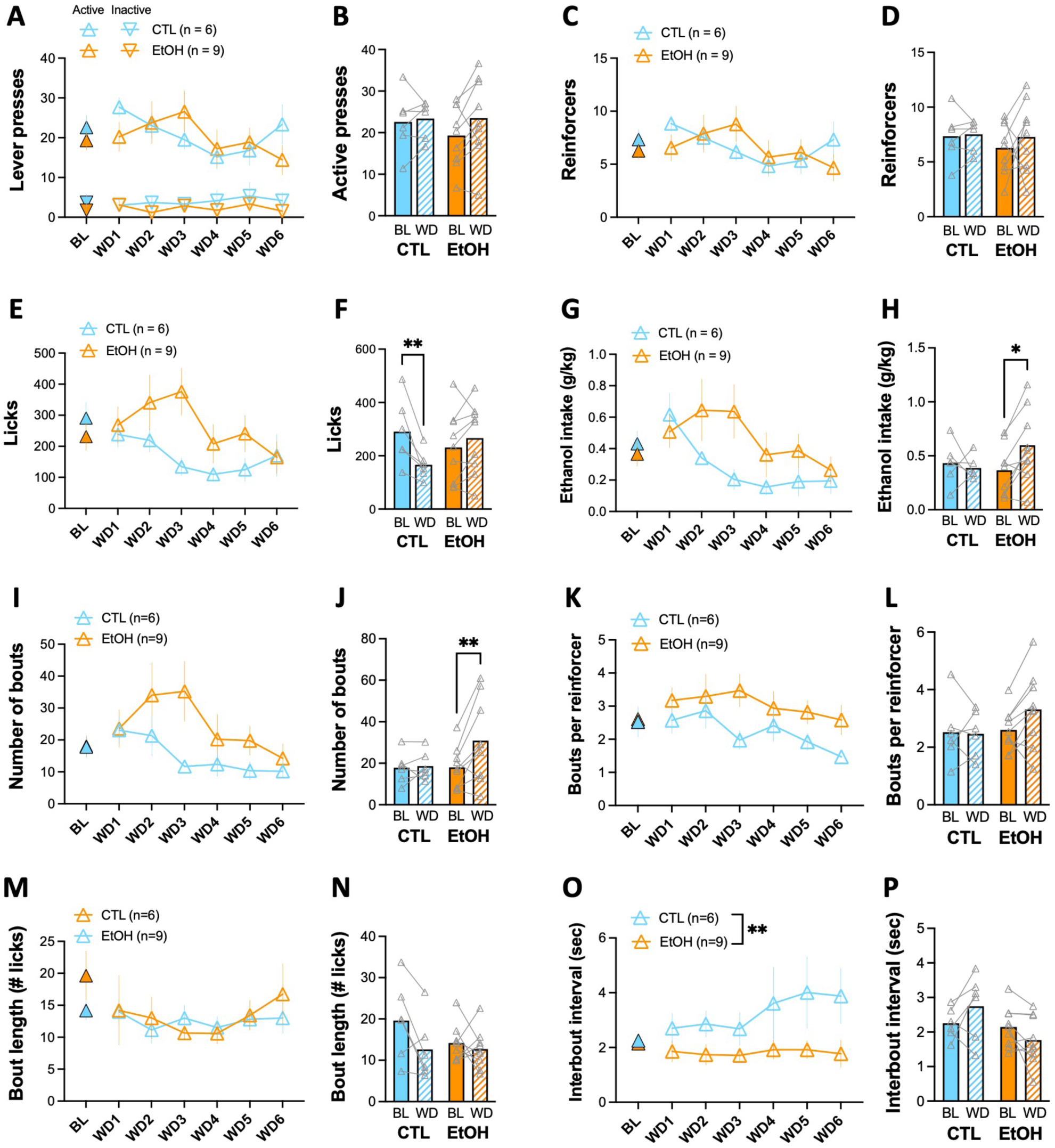
Transient withdrawal-induced escalation of ethanol intake in female Drinkers chronically exposed to ethanol liquid diet. Number of active lever presses **(A)** and reinforcers earned **(C)** is unchanged after chronic liquid diet exposure. A non-significant increase in consumption as indicated by licks **(E)** and intake **(G)** is observed in Drinkers exposed to ethanol, but not control, liquid diet. This is associated with a similar non-significant increase in the total number of drinking bouts **(I)** during a session and number of bouts per reinforcer access period **(K)**. Bout length **(M)** is similar between groups across time. Interbout interval **(O)** is significantly greater in control than ethanol liquid diet-exposed rats. Comparison of performance during baseline and the first three withdrawal sessions confirms the absence of change in appetitive responding **(B, D)** and reveals a significant decrease in licks **(F)** in control diet exposed Drinkers and an increase in intake **(H)** in ethanol diet exposed Drinkers. This is associated with a significant increase in the total number of drinking bouts **(J)** in ethanol exposed Drinkers relative to controls in the absence of any differences in number of bouts per reinforcer access period **(L)**, bout length **(N)**, or interbout interval **(P)**. *p<0.05; **p<0.01

Despite the absence of robust significant differences in operant performance during repeated acute withdrawal sessions, visual inspection of the data revealed a separation in consummatory behaviors between female rats exposed to control and ethanol liquid diet that occurred during the first three withdrawal sessions relative to the last three sessions. To examine this more directly, we next compared average baseline operant performance with performance during the first three acute withdrawal test sessions. Similar to our observations above, a two-way RM ANOVA examining changes in active lever pressing did not find significant main effects of either liquid diet [F (1, 13) = 0.1664; p=0.6900] or time [F (1, 13) = 3.036; p=0.1050] or a significant diet by time interaction [F (1, 13) = 1.375; p=0.2620] (**Figure 6B**). No significant main effects of diet [F (1, 13) = 0.1073; p=0.7485] or time [F (1, 13) = 3.076; p=0.1030] or diet by time interaction [F (1, 13) = 1.951; p=0.1859] were observed for reinforcers earned either (**Figure 6D**). In contrast, while no main effect of liquid diet [F (1, 13) = 0.1154; p=0.7395] was observed with regard to number of licks during the operant session, a two-way RM ANOVA uncovered a significant main effect of time [F (1, 13) = 4.615; p=0.0511] as well as a significant interaction between diet and time [F (1, 13) = 14.69; p=0.0021]. Although the average number of licks appeared to increase slightly during withdrawal sessions in rats exposed to ethanol liquid diet, post-hoc comparisons using a Sidak correction unexpectedly found that licks were significantly lower in the first three acute withdrawal sessions relative to baseline in female rats exposed to control, but not ethanol, liquid diet (p=0.0039; **Figure 6F**). A two-way RM ANOVA examining differences in intake also uncovered a significant diet by time interaction [F (1, 13) = 5.517; p=0.0353] but no main effects of either factor [diet: F (1, 13) = 0.3921; p=0.5420; time: F (1, 13) = 2.535; p=0.1353]. In contrast to licks, post-hoc comparisons of this interaction using a Sidak correction found that rats exposed to ethanol liquid diet had significantly greater intake during the first three acute withdrawal sessions compared to baseline (p=0.0163), whereas intake in rats exposed to control diet was unchanged (p=0.8656; **Figure 6H**).

To explore the basis of the observed withdrawal-induced increase in ethanol intake in ethanol liquid diet exposed rats in greater depth, we next investigated changes in drinking microstructure. Similar to our analysis of licks and overall intake, two-way RM ANOVAs did not uncover any significant main effects or interactions when comparing the average number of drinking bouts (**Figure 6I**), bouts per access to reinforcer (**Figure 6K**), or bout length (**Figure 6M**) before and after chronic liquid diet and between diet groups (all p values > 0.05). The same analysis of interbout interval revealed a significant main effect of diet [F (1, 13) = 9.956; p=0.0076] with rats that received control diet taking more time between drinking bouts than rats that received ethanol diet (**Figure 6O**). Although visually, this effect appears to be primarily driven by an increase in interbout interval after diet exposure, no significant main effect of time [F (2.937, 38.18) = 0.7248; p=0.5407] or time by diet interaction [F (6, 78) = 0.8750; p=0.5172] was observed. Comparisons restricted to the first three days of acute withdrawal testing, when a significant increase in overall intake was observed in rats exposed to ethanol liquid diet, revealed a significant withdrawal-induced increase in the number of drinking bouts (**Figure 6J**) in rats that received ethanol (p=0.0073), but not control, liquid diet. This was evident using Sidak’s multiple comparisons after uncovering a significant main effect of time [F (1, 13) = 5.693; p=0.0329], which occurred in the absence of significant effects of diet [F (1, 13) = 0.9811; p=0.3400] or time by diet interaction [F (1, 13) = 4.348; p=0.0573]. No significant differences were observed for number of drinking bouts per reinforcer access (all p values > 0.05; **Figure 6L**). Analysis of bout length revealed a significant main effect of time [F (1, 13) = 5.448; p=0.0363] with rats exhibiting an overall decrease in bout length during acute withdrawal (**Figure 6N**). Although this effect appears to be driven by a decrease in bout length in rats exposed to control diet, this analysis did not uncover a significant main effect of diet [F (1, 13) = 0.8910; p=0.3624] or time by diet interaction [F (1, 13) = 2.326; p=0.1512]. Finally, analysis of interbout interval (**Figure 6P**) found a significant time by diet interaction [F (1, 13) = 5.432; p=0.0365] in the absence of main effects of time [F (1, 13) = 0.09202; p=0.07664] or diet [F (1, 13) = 3.072; p=0.1032]. However, multiple comparisons using Sidak’s correction found no significant change in interbout interval from baseline in either control (p=0.2130) or ethanol (p=0.2483) exposed rats. Altogether, these data indicate that consummatory behavior increases during acute withdrawal in rats exposed to chronic ethanol liquid diet, at least transiently during repeated testing, in the absence of any changes in appetitive responding. This increase is driven primarily by an increase in the number of drinking bouts in the absence of changes to bout length or interbout interval.

### 3.6 Operant performance does not change in Responders during acute withdrawal from chronic ethanol liquid diet

Because our prior work [17] uncovered withdrawal-induced changes in operant performance in male rats of the Responder phenotype, we subsequently performed the same analysis of operant behavior in female Responders following control and ethanol liquid diet. Two-way RM ANOVAs of active lever presses and reinforcers earned found no significant main effects of either diet [lever press: F (1, 23) = 0.1920; p=0.6653; reinforcer: F (1, 23) = 0.1310; p=0.7207] or time [lever press: F (5.027, 115.6) = 1.494; p=0.1969; reinforcer: F (5.139, 118.2) = 1.406; p=0.2258] but did uncover significant diet by time interactions [lever press: F (6, 138) = 3.334; p=0.0043; reinforcer: F (6, 138) = 3.468; p=0.0032] for both variables (**Figure 7A,C**). Despite this, post-hoc comparisons did not find any significant between group differences in these measures during any given operant session (all p values > 0.20) suggesting that differences in lever pressing and reinforcers earned between rats exposed to control and ethanol liquid diet were minimal. A two-way RM ANOVA examining change in licks found no significant main effects [diet: F (1, 23) = 0.01694; p=0.8975; time: F (1.506, 34.63) = 0.8167; p=0.4190] or time by diet interaction [F (6, 138) = 0.9698; p=0.4481] (**Figure 7E**). The same analysis for ethanol intake found no significant effect of diet [F (1, 23) = 0.6057; p=0.4443] but did reveal a significant main effect of time [F (2.860, 65.78) = 3.419; p=0.0240] in the absence of a significant interaction between diet and time [F (6, 138) = 1.010; p=0.4213]. Post-hoc comparisons using a Dunnett correction to explore the main effect of time found that ethanol intake during the first acute withdrawal session was significantly greater in both groups compared to intake at baseline (p=0.0420; **Figure 7G**). Although statistically significant, the relevance of this difference is questionable given the very low levels of average ethanol intake observed at both baseline (0.03 ± 0.004 g/kg) and during the first withdrawal session (0.07 ± 0.012 g/kg).

**Figure 7.**
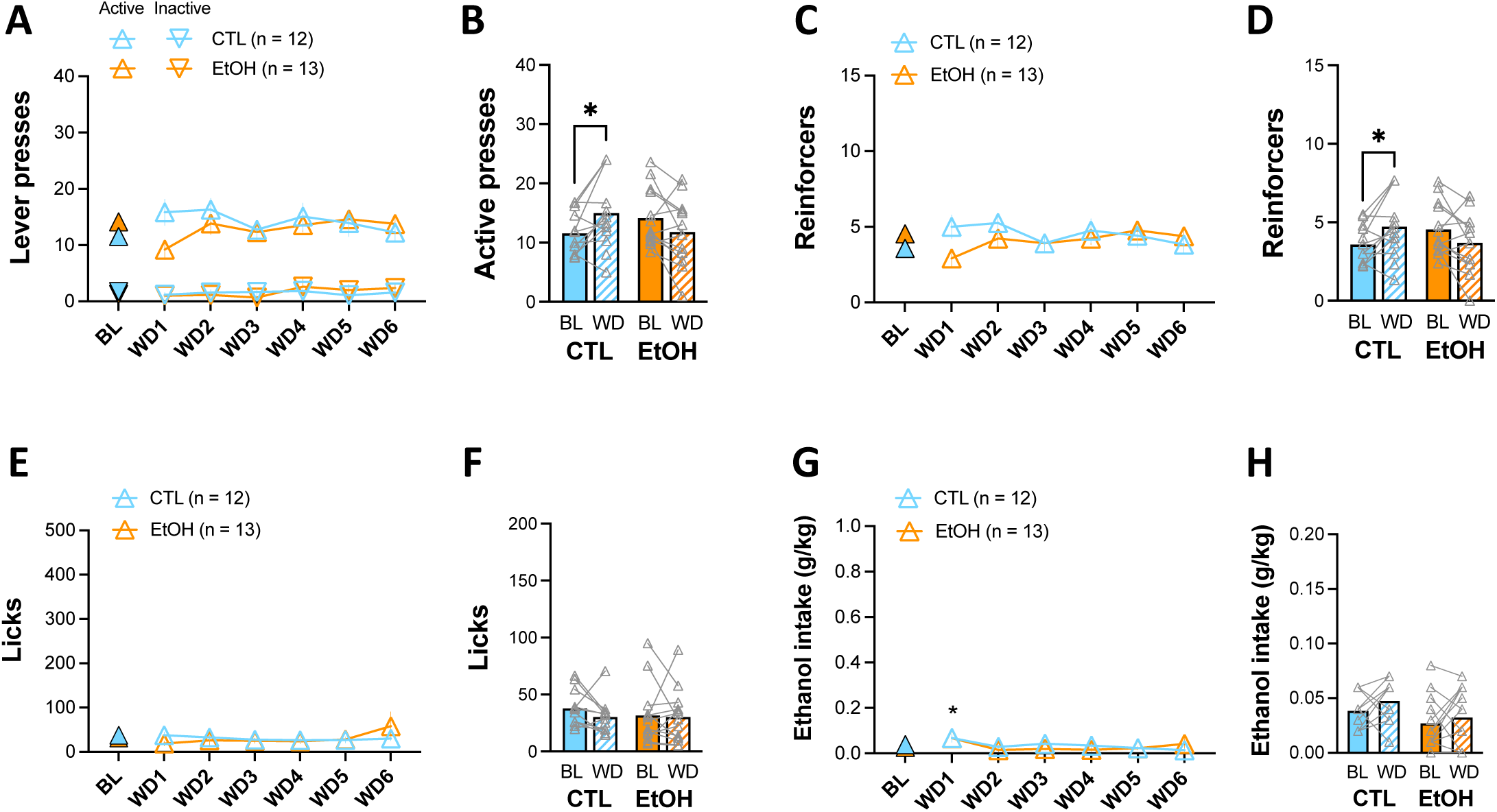
Minimal effect of liquid diet exposure on appetitive and consummatory behaviors in female Responders. When examined across the 6 acute withdrawal testing sessions, Responders showed no significant change in the number of active lever presses **(A),** reinforcers earned **(C)**, or licks **(E)** after chronic liquid diet exposure. A modest, but statistically significant, increase in ethanol intake **(G)** was observed in all Responder rats during the first withdrawal test session relative to baseline independent of liquid diet group. However, this effect was transient as intake was not significantly different from baseline on subsequent test days. Comparison of performance during baseline and the first three withdrawal sessions revealed a significant increase in active lever presses **(B)** and reinforcer access **(D)** during withdrawal in rats exposed to control, but not ethanol, diet. No differences in either licks **(F)** or intake **(H)** were observed. *p<0.05

Given that we observed between-group separation, albeit modest, in appetitive performance during early withdrawal sessions in Responders, we proceeded with the same analyses we applied to Drinkers by examining operant performance restricted to the first three withdrawal test sessions. Two-way RM ANOVAs of lever presses during baseline and the first three withdrawal sessions revealed a significant diet by time interaction [F (1, 23) = 9.688; p=0.0049] in the absence of main effects [time: F (1, 23) = 0.3251; p=0.5741; diet: F (1, 23) = 0.02383; p=0.8787]. Post-hoc comparisons using a Sidak correction showed that lever pressing was significantly increased during withdrawal from baseline performance in female rats that received control, but not ethanol, liquid diet (p=0.0352; **Figure 7B**). Similar results were uncovered when examining number of reinforcers earned. Again, a two-way RM ANOVA found a significant interaction between liquid diet exposure and time [F (1, 23) = 9.998; p=0.0044] in the absence of main effects for either factor [diet: F (1, 23) = 0.003094; p=0.9561; time: F (1, 23) = 0.2244; p=0.6402]. Similar to our observations with active lever presses, post-hoc comparisons using a Sidak correction revealed that female rats given control liquid diet earned significantly more reinforcers during early withdrawal sessions compared to during baseline testing (p=0.0378), whereas the number of reinforcers earned in ethanol diet exposed rats was unchanged (**Figure 7D**). In contrast, measures of consummatory behavior were unchanged in rats exposed to either control or ethanol liquid diet. This was evident in two-way RM ANOVAs, which found no significant main effects or interactions for either licks [time: F (1, 23) = 0.9728; p=0.3343; diet: F (1, 23) = 0.1961; p=0.6620; time x diet: F (1, 23) = 0.4471; p=0.5104] or intake [time: F (1, 23) = 1.874; p=0.1842; diet: F (1, 23) = 3.682; p=0.0675; time x diet: F (1, 23) = 0.1266; p=0.7252] (**Figure 7F,H**). Collectively, these data reveal a slight, but statistically significant, increase in appetitive responding in female Responders that received control diet that is absent in Responders that received ethanol liquid diet.

## 4. Discussion

Results from the present study closely parallel our previous observations in male rats [17] by revealing in female rats: 1) the presence of the same three distinct operant phenotypes, 2) that the prevalence of each phenotype differs across strains, and 3) that the Responder phenotype would be undetected in the absence of direct measures of consumption during operant testing. Our results further show that operant performance cannot be determined in female rats from average ethanol intake across three weeks of home cage drinking. Finally, although female Drinkers exhibited significant withdrawal-induced escalation of operant ethanol self-administration following chronic ethanol liquid diet exposure, this escalation was limited to consummatory behaviors and transient, returning back to baseline levels after several test sessions.

A number of studies have observed greater ethanol intake in female than male rats of various strains during home cage drinking (e.g., [32–34]). Unlike these studies, average levels of daily intake at the end of the two-bottle choice period in the current study were similar to levels observed in our previous work in males [17]. However, the increase in ethanol intake that is regularly observed in males across the first few drinking sessions [17, 18, 28] was not present in the current study. Instead, females of all three strains exhibited relatively consistent levels of daily ethanol intake from the first to the last drinking session. Thus, females appear to have higher levels of ethanol intake than males during early, but not later, two-bottle choice access. Whether this phenomenon is routinely observed in other studies is somewhat difficult to determine as many studies collapse data across days or weeks. Nevertheless, some work suggests that this may be a common pattern of home cage drinking in female rats. For example, similar to our findings, previous work in alcohol-preferring P rats, which directly compared drinking across sexes, found greater ethanol intake in females than males early during home cage drinking but similar levels of intake between sexes after the first 2-3 weeks of drinking [12]. In addition, despite concluding that intake is greater in females than males, data from Morales et al. [33] show that the gap in intake between the sexes narrows over increasing weeks of home cage access. Interestingly, rats from all three strains in the current study exhibited similar levels of intake throughout the home cage drinking phase of the study. This is unlike our previous work in males, which found that Long Evans rats drink significantly more than Wistar and Sprague Dawley strains [17]. Although many studies have examined strain differences in ethanol drinking, these have been performed almost exclusively in males. Thus, these data provide new, and much needed, insight into potential strain and sex differences (or lack thereof) in home cage ethanol drinking.

Similar to our previous work, all three operant phenotypes were evident in females of all three rat strains. In addition, the prevalence of each phenotype in females aligned closely with our observations in males with the Drinker phenotype being most prevalent in the Long Evans strain and least prevalent in the Sprague Dawley strain. In agreement with our earlier work, Responders were highly prevalent in female Wistar and Sprague Dawley strains [17]. The striking similarities between studies suggests that the prevalence of each operant phenotype may be a characteristic of each strain independent of sex.

Comparable to our findings in males, baseline operant performance was not different between strains of female rats within a given phenotype. In addition, average baseline ethanol intake (g/kg) in female Drinkers paralleled our previous observations in males (∼0.5 g/kg/session) [17]. However, females had fewer active lever presses and reinforcer deliveries as well as fewer licks per session compared to what we observed previously in males. This is not surprising given that less total volume is required for females, which have smaller average body weight, to consume an equivalent dose of ethanol to males. Although not all studies agree (e.g., [35]), our data is supported by a handful of other studies that show similar rates of ethanol intake between male and female rats during operant ethanol self-administration [12, 20, 30, 36].

Baseline performance in female Responders also exhibited striking similarities to baseline performance documented in males in our previous work [17]. Indeed, female Responders exhibited a similar degree of appetitive responding (with clear discrimination between the active and inactive levers) accompanied by a similar absence of consumption, as was observed in males [17]. The disconnect between appetitive and consummatory behavior in Responders was further confirmed by the absence of a significant correlation between behaviors. The significant discrepancy in Responders between actual intake as measured by change in bottle weight and intake extrapolated from an estimate of volume consumed per reinforcer delivery calls attention to the risk posed by inferring consumption based on appetitive responding. These data support previous studies underscoring the importance of measuring both appetitive and consummatory behaviors during operant ethanol self-administration.

In females, home cage drinking was indistinguishable across phenotypes. Thus, future Drinkers, Responders, and Nonresponders all exhibited similar average levels of intake and no significant relationship was observed between home cage and operant drinking. Previously in males, we found that future Drinkers consumed significantly more ethanol on average during home cage drinking than future Responders and Nonresponders. However, there was significant overlap in average daily intake across the entire population of each phenotype and, similar to our observation in females, intake during home cage and operant sessions was not significantly correlated [17]. Together, these data indicate that consumption after three weeks of home cage drinking cannot predict operant performance in either male or female rats. However, intake after a longer period of home cage drinking may be informative [37].

Withdrawal-induced escalation of ethanol intake is frequently used to model uncontrolled drinking, which is a primary feature of AUD [38]. However, few studies have examined this phenomenon in female rats. Previous work measuring escalation using a home cage drinking paradigm suggested that, unlike males, a ceiling effect may contribute to the absence of escalation in females [33]. To our knowledge, only two studies have examined escalation in females using an operant paradigm. Both of these studies report significant withdrawal-induced escalation of operant ethanol self-administration in female Wistar rats [20, 30], but not female Long Evans rats [20]. Similar to males, female Drinkers in the current study exhibited significant withdrawal-induced escalation of operant ethanol self-administration after chronic ethanol liquid diet exposure. Interestingly, while acute withdrawal was associated with a significant increase in the number of drinking bouts in addition to greater overall intake, a concomitant increase in active lever presses or reinforcer delivery was not observed. This is in contrast to our prior observations in male Drinkers who exhibited a significant withdrawal-induced increase in both appetitive and consummatory responses [17]. This suggests that, unlike males, females drank more ethanol during acute withdrawal without increasing the amount of effort required. How these data compare to the two prior studies in females is difficult to determine as neither study reports appetitive responding, instead reporting intake (g/kg) only [20, 30]. While the current study was not sufficiently powered to examine potential strain differences in escalation of ethanol intake, it should be noted that, similar to baseline, operant performance was comparable across strains during acute withdrawal testing (not shown).

Curiously, we found that while female Drinkers exhibit increased consumption during acute withdrawal relative to baseline, this effect was transient in nature lasting for only the first three of six acute withdrawal test sessions. This is unlike our previous observations in males who exhibited persistent escalation over nine separate acute withdrawal test sessions [17]. Prior work is unable to provide insight into whether the transient escalation observed in the current study is a common feature in female rats. Matzeu et al. [30] reported escalation in intake across three acute withdrawal sessions, corresponding to the same time frame during which we also observe escalation. Although somewhat unclear, Priddy et al. [20] appear to have measured intake over many withdrawal sessions. However, data showing escalation are collapsed across sessions obscuring the ability to appreciate whether operant performance varied over time. While the basis for the transient escalation observed in the current study is unclear, it is possible that females experience tolerance to symptoms of withdrawal over repeated acute withdrawal episodes. Although this remains conjecture in the absence of data supporting this notion, it is conceivable that the loss of escalation observed after the first three test sessions could be driven by attenuation of withdrawal symptoms. Future studies designed to replicate the current findings in tandem with measures of withdrawal symptoms will be crucial to develop stronger conclusions regarding this possibility.

Behavioral performance was largely unaffected by liquid diet exposure in Responders. This is contrary to our previous observations, which found that male Responders increased appetitive behavior following ethanol liquid diet exposure and that a small subset of these rats engaged in substantive drinking behavior [17]. In contrast, in the current study none of the female Responders exhibited changes in behavior indicative of a “switch” in operant phenotype after chronic ethanol exposure. This suggests that the capacity to transition from one phenotype to another may be restricted to males and/or occur only rarely.

In conclusion, although small differences exist, by showing that female rats exhibit the same three operant phenotypes and exhibit similar operant performance at baseline, the present study largely replicates prior work performed in males. It extends these data further by revealing that escalation of ethanol intake during acute withdrawal is transient in females and limited to consummatory behaviors while effort required to acquire access to the reinforcer is unchanged. These data provide new insight into potential sex and strain differences in home cage and operant ethanol drinking behavior. Moreover, our findings emphasize the crucial need for inclusion of both male and female subjects in preclinical studies of AUD.

## Acknowledgements

Thanks to Alex Brown, Shree Srinivasan, Nidhi Chetan, Arshdeep Bal, and E. Margaret Starr for technical assistance. This work was supported by the National Institute on Alcohol Abuse and Alcoholism at the National Institutes of Health (P50 AA022538 and R01 AA029130 to EJG).

